# A Novel Organoid Model of In Vitro Spermatogenesis Using Human Induced Pluripotent Stem Cells

**DOI:** 10.1101/2021.06.04.447122

**Authors:** Meghan Robinson, Luke Witherspoon, Stephanie Willerth, Ryan Flannigan

## Abstract

Infertility is thought to be caused by genetic mutations and dysfunction in the cellular niche where spermatogenesis takes place. An understanding of the specialized cellular processes which drive spermatogenesis is needed to develop treatments; however, the development of *in vitro* systems to study these cells has been hindered by our reliance on rarely available human testicular tissues for research. Human induced pluripotent stem cells (hiPSCs) can be used to derive human testicular-like cells, and thus provide an avenue for the development of *in vitro* testicular model systems. Therefore, this study set out to engineer a human testicular tissue model using hiPSCs for the first time. We demonstrate the ability of hiPSC-derived testicular cells to self-organize and mature into testicular-like tissues using organoid culture. Moreover, we show that hiPSC-derived testicular organoids promote testicular somatic cell maturation and spermatogenesis up to the post-meiotic spermatid stage. These hiPSC-derived testicular organoids have the potential to replace rarely available primary testicular tissues to further infertility research in an *in vitro* setting.

## Introduction

The generation of sperm is a complex process dependent upon the relationships between multiple cell types within the testis.[1] Pinpointing the causes of infertility therefore requires an understanding of the cellular niche where sperm are generated. Advances in single cell sequencing of testicular tissue have begun to shed light on the transcriptomic signatures of the cell types within the niche,[2-8] however the development of *in vitro* tools to study their interactions and functionalities remains elusive.[9]

Historically, animal models have been the standard in modeling human diseases, however there are specifically human biological processes such as infertility whose complexity and inter-individual variability cannot be accurately modeled using animals.[2, 3, 6, 10-15] Consequently, efforts to model infertility have turned to the development of human *in vitro* systems. One such system that is proving to be a particularly valuable tool for drug discovery, toxicology and personalized medicine is the organoid system.[16] Organoids are 3-dimensional cellular systems that are histologically similar to native tissues and organs, and can recapitulate many of their multicellular processes.[10, 17, 18] They can be generated from pluripotent or multipotent stem cells, and are patterned via the same innate pathways as embryogenesis to form miniature functional units of a larger tissue or organ. Efforts to develop human testicular organoid systems have been significant.[19-24] However, despite success in animal testicular organoid systems, determination of the *in vitro* conditions necessary for human testicular organoid generation remains a challenge, compounded by the rare accessibility of human testicular tissue for research.[24]

A potential solution is to engineer testicular tissues using human induced pluripotent stem cells (hiPSCs), a Nobel Prize-winning technology pioneered by Shinya Yamanaka in 2007[25] that has since revolutionized *in vitro* modeling and personalized medicine.[26] hiPSCs are adult cells reprogrammed into pluripotent stem cells through the forced expression of a set of key transcription factors. Like embryonic stem cells, they can give rise to all tissues of the body and regenerate indefinitely, providing an inexhaustible resource for *in vitro* modeling. Moreover, unlike embryonic stem cells, they can be sourced from a variety of adult cell types, paving the way for personalized medicine.

To date the use of hiPSCs to generate testicular organoids has yet to be investigated, likely because methods for the derivation of the major cellular components of the testicular niche from hiPSCs have only recently been established.[27-30] Using these derivation strategies, we engineered and evaluated a novel model of hiPSC-derived testicular organoids. We assessed their ability to mature and re-organize into testicular-like tissues and support spermatogenesis using immunohistochemical staining techniques and real time quantitative polymerase chain reaction (RT-qPCR) assays. This work reports for the first time the generation of testicular-like tissues from hiPSCs, and demonstrates their potential to model spermatogenesis.

## Results

### Cellular identity characterization

As a first step towards generating testicular organoids we derived the main components of the testicular niche from hiPSCs: Sertoli cells, the main drivers of spermatogenesis; Leydig cells, the producers of testosterone required for Sertoli cell function; endothelial cells, a source of growth factors vital to the homeostasis of the niche; peritubular myoid cells, the contractile smooth muscle cells which facilitate transport of spermatozoa to the epididymis; and spermatogonial stem cells (SSCs), the self-renewing stem cell pool that gives rise to spermatozoa (Figure 1A). The acquisition of testis-specific phenotypes in the hiPSC-derived cells was confirmed by immunocytochemistry staining for cell-specific protein expression signatures and markers of maturation (Figure 1B).

**Figure 1.**
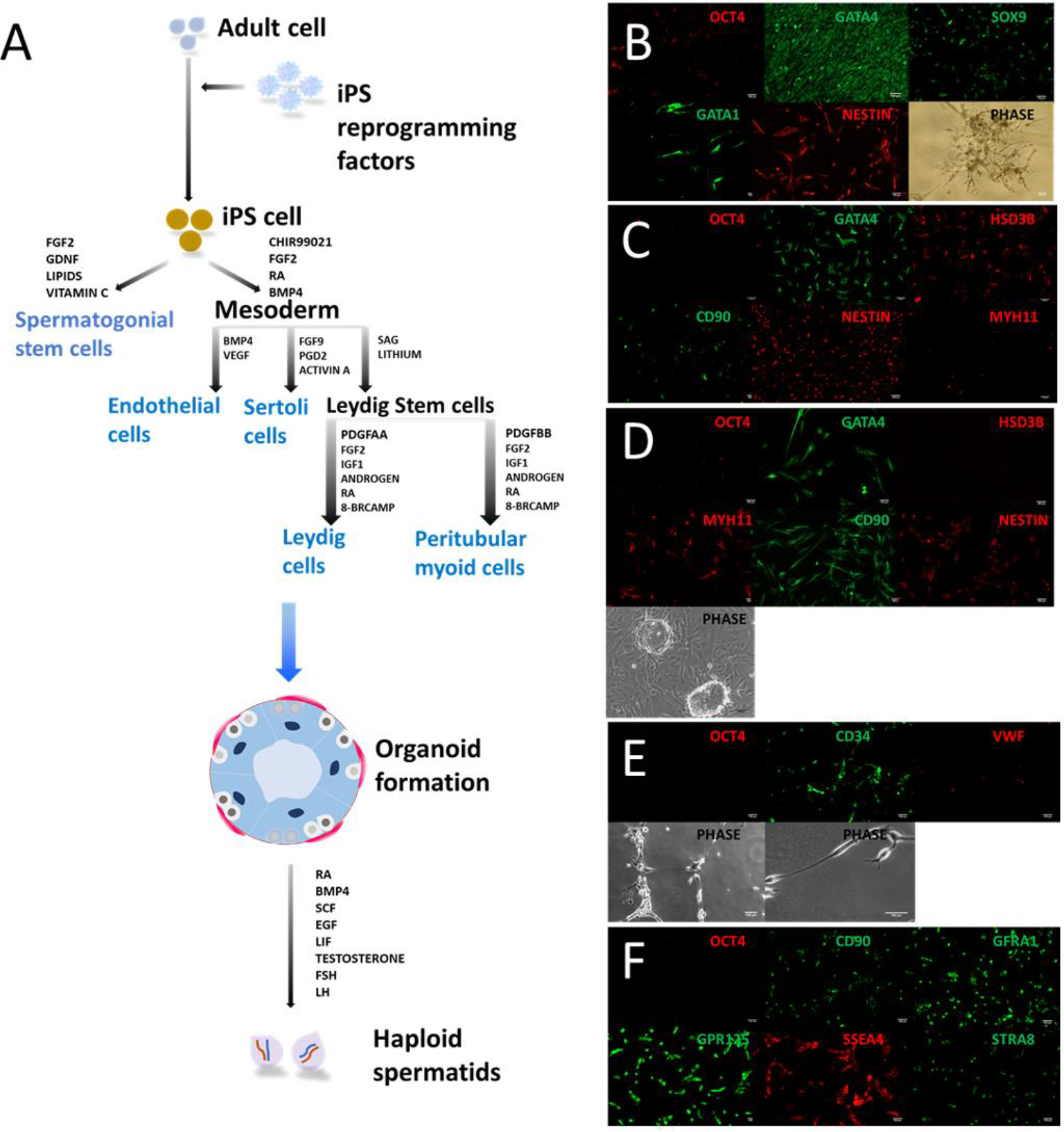
hiPSC derivation of testicular cell types. **A)** Chart of the derivation of testicular cells from hiPSCs and organoid culture conditions. **B)** Immunocytochemistry staining of hiPSC-derived Sertoli cells, and a phase contrast image showing cord formation in 3-D culture. **C)** Immunocytochemistry staining of hiPSC-derived Leydig cells. **D)** Immunocytochemistry staining of hiPSC-derived peritubular myoid cells, and a phase contrast image showing ring formation in 2-D culture. **E)** Immunocytochemistry staining of hiPSC-derived endothelial cells, and phase contrast images of tubulogenesis in low and high magnifications. **F)** Immunocytochemistry staining of hiPSC-derived spermatogonial stem cells. All scale bars are 100 µm. **Abbreviations:** 8-BRCAMP = 8-bromo-cAMP, BMP4 = Bone Morphogenic Protein 4, CD34 = CD34 Antigen, CD90 = Thy-1 Cell Surface Antigen, EGF = Epidermal Growth Factor, FGF2 = Fibroblast Growth Factor 2, FGF9 = Fibroblast Growth Factor 9, FSH = Follicle Stimulating Hormone, GATA1 = GATA-Binding Protein 1, GATA4 = GATA-Binding Protein 4, GDNF = Glial Cell-Derived Neurotrophic Factor, GFRA1 = GDNF Family Receptor Alpha 1, GPR125 = G-Protein Coupled Receptor 125, HSD3B = Hydroxy-Delta-5-Steroid Dehydrogenase 3 Beta- And Steroid Delta-Isomerase 1, IGF1 = Insulin-Like Growth Factor 1, iPS = induced pluripotent stem, LH = Luteinizing Hormone, LIF = Leukemia Inhibitory Factor, MYH11 = Myosin Heavy Chain 11, OCT4 = Octamer Binding Protein 4, PDGFAA = Platelet-Derived Growth Factor-AA, PDGFBB = Platelet-Derived Growth Factor-BB, PGD2 = Prostaglandin D2, RA = retinoic acid, SAG = smoothened agonist, SCF = Stem Cell Factor, SOX9 = SRY-Box Transcription Factor 9, SSEA4 = Stage-Specific Embryonic Antigen 4, STRA8 = Stimulated By Retinoic Acid 8, VEGF = Vascular Endothelial Growth Factor, VWF = Von Willebrand Factor.

hiPSC-derived Sertoli cells (hSCs) expressed SOX9, a Sertoli cell-specific transcription factor,[31, 32] and GATA4, a testis-specific transcription factor,[33, 34] confirming a Sertoli cell phenotype (Figure 1B). Loss of the pluripotency factor OCT4[35] further confirmed a differentiated status (Figure 1B). However, expression of Nestin, a cytoskeletal filament found in differentiating cells,[36] indicated a state of immaturity (Figure 1B). Confirming this, only a small number of hSCs expressed GATA1, a marker of mature Sertoli cells (Figure 1B).[37] *In vivo*, immature Sertoli cells form cords which mature during puberty into the seminiferous tubules of the testis,[38] therefore we tested the hSCs for cord-forming functionality by aggregating and seeding them on a 3-dimensional layer of Matrigel[39]. Cords were seen to develop within the Matrigel layer within 24 hours, and became increasingly elongated and branched over the course of 7 days, in keeping with an immature Sertoli cell phenotype (Figure 1B).

hiPSC-derived Leydig Cells (hLCs) were negative for the pluripotency factor OCT4, and positive for the testicular marker GATA4 and the differentiation marker Nestin, in keeping with an immature testicular phenotype (Figure 1C). They were also positive for HSD3B, a Leydig cell steroidogenic enzyme required for testosterone synthesis (Figure 1C), confirming Leydig cell character.[40] The mature Leydig cell marker INSL3[40] was negative, while CD90, a Leydig progenitor marker[40, 41], was positive, indicating an immature state (Figure 1C).

hiPSC-derived peritubular myoid cells (hPTMs) were likewise negative for the hiPSC marker OCT4, and positive for the testicular marker GATA4, the progenitor marker CD90, and the differentiation marker Nestin (Figure 1D). Furthermore, hPTMs were positive for peritubular myoid cell marker MYH11[42], confirming their peritubular myoid cell character (Figure 1D). hPTMs were also noted to form ring-like structures in culture, a previously documented characteristic of purified peritubular myoid cell cultures (Figure 1D).[43]

hiPSC-derived endothelial cell (hEC) OCT4 expression was negative as expected, and the endothelial progenitor marker CD34 was positive,[44] confirming vascular differentiation (Figure 1E). VWF, an endothelial cell protein necessary for hemostasis, was only rarely observed by staining, suggesting that the majority of the cells maintained an immature phenotype (Figure 1E).[45] A critical function of endothelial cells is blood vessel formation,[46] so we tested the ability of the hECs to undergo tubulogenesis. After 24 hours in culture on a 3-dimensional layer of Matrigel, hECs formed branching networks of tubular structures as expected (Figure 1E). Immunocytochemistry analyses of the hiPSC-derived SSCs (hSSCs) confirmed positive expression of several well-known SSC markers, including CD90, GFRA1, SSEA4, and GPR125,[2, 47, 48] and negative expression for the hiPSC pluripotency factor OCT4 (Figure 1F). Expression of the pre-meiotic stage SSC gene STRA8[49, 50] was also noted (Figure 1F), in keeping with reports of primary SSCs grown in monolayer culture.[20, 21, 51]

### hiPSC-derived testicular organoid characterization

hiPSC-derived testicular cells self-assembled into organoids when cultured overnight in AggreWell™800 microwell plates (Figure 2I), and were transferred to non-adherent plates for suspension culture for 12 days (Figure 2J). The culture medium was designed to mimic *in vivo* conditions through supplementation with hormones FSH, LH and testosterone, to stimulate somatic cell function and maturation,[52, 53] and the Sertoli cell-secreted factors BMP4, SCF and retinoic acid, as studies have shown that these factors are required for SSC entry into differentiation.[50, 54-59] The factors EGF and LIF were added for their known effect on promoting viability of SSCs *in vitro*,[60] and because differentiating germ cells and somatic cells express receptors for these factors *in vivo*, although reasons for their expression are not well understood.[61-64]

**Figure 2.**
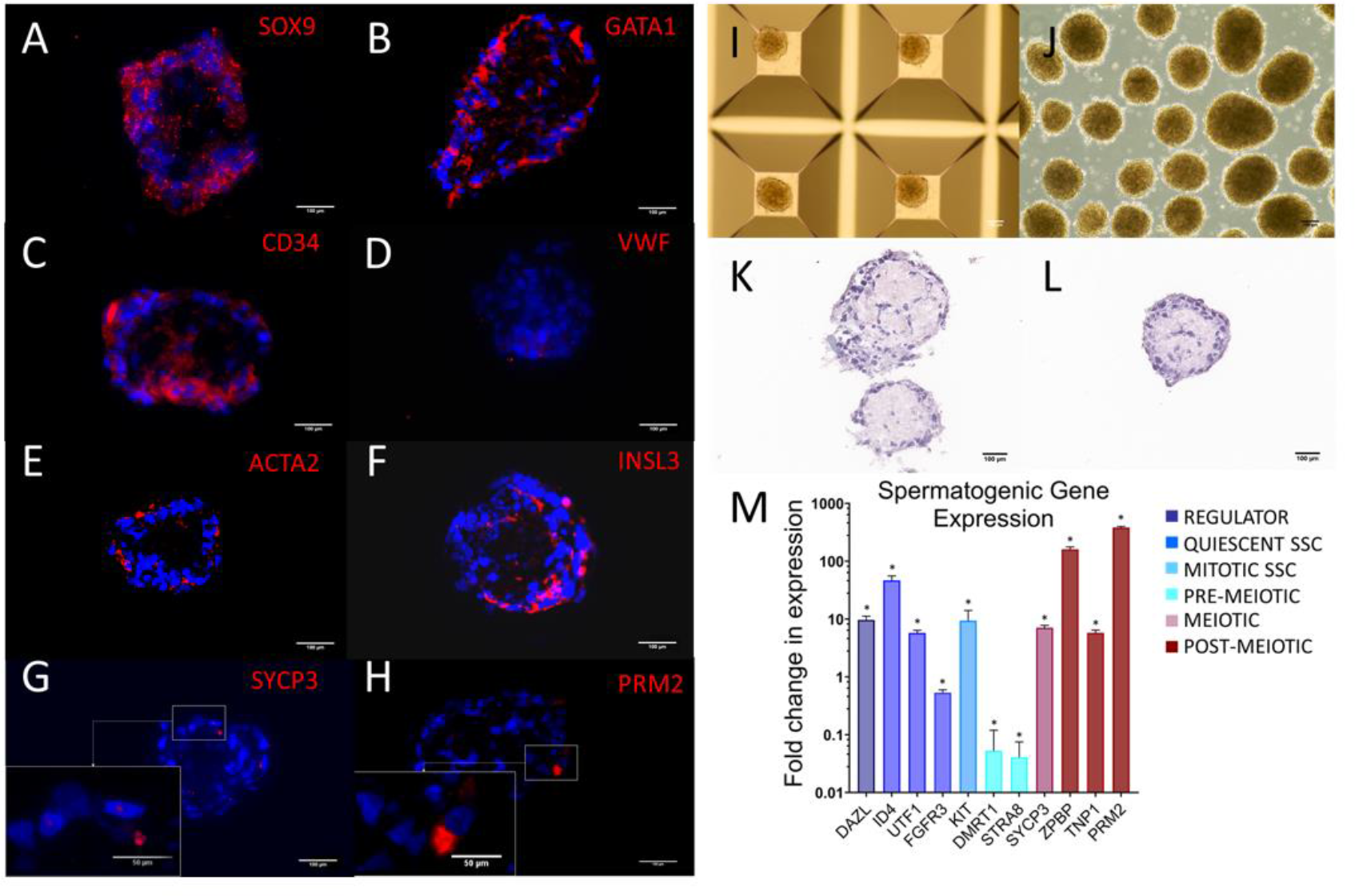
hiPSC-derived testicular organoids. **A-H)** Immunostaining of organoid sections showing their internal organization and maturation: SOX9^+^/GATA1^+^ = mature Sertoli cells, ACTA2^+^ = peritubular myoid cells, INSL3^+^ = mature Leydig cells, CD34^+^/VWF^-^ = immature endothelial cells, SYCP3^+^ = meiotic germ cells, PRM2^+^ = post-meiotic germ cells. **I)** Phase image of organoids 24 hours after formation in microwells. **J)** Phase image of organoids in suspension culture. **K-L)** H&E-stained organoid sections showing the cellular structures. **M)** Fold change in spermatogenic gene expression of germ cells in the organoids after 12 days (compared to day 0): DAZL is a regulatory gene involved SSC maintenance and differentiation, UTF1, ID4 and FGFR3 are markers of quiescent SSCs, KIT is a marker of mitotic differentiating SSCs, DMRT1 and STRA8 are pre-meiotic markers of early differentiating germ cells, SYCP3 is a marker of meiotic differentiating germ cells, ZPBP, TNP1 and PRM2 are markers of post-meiotic germ cells undergoing spermiogenesis. All scale bars are 100 µm except the high magnification SYCP3 and PRM2 images where they are 50 µm. **Abbreviations:** ACTA2 = Actin Alpha 2, Smooth Muscle, INSL3 = Insulin-Like 3, CD34 = CD34 Molecule, DAZL = Deleted in Azoospermia-Like, DMRT1 = Doublesex And Mab-3 Related Transcription Factor 1, FGFR3 = Fibroblast Growth Factor Receptor 3, GATA1 = GATA-Binding Protein 1, H&E = Hematoxylin and Eosin, ID4 = Inhibitor Of DNA Binding 4 HLH Protein, INSL3 = Insulin Like 3, KIT = KIT Proto-Oncogene Receptor Tyrosine Kinase, PRM2 = Protamine 2, SOX9 = SRY-Box Transcription Factor 9, SSC = spermatogonial stem cell, STRA8 = Stimulated By Retinoic Acid 8, SYCP3 = Synaptonemal Complex Protein 3, TNP1 = Transition Protein 1, VWF = Von Willebrand Factor, ZPBP -= Zona Pellucida Binding Protein.

After 12 days in culture, organoids were harvested for gene expression and histochemical analyses. Viability staining was not possible due to high levels of autofluorescence, however the organoids maintained a healthy morphology characterized by a well-defined, bright outer boundary and no visibly dark areas of necrosis (Figure 2J).

RT-qPCR analysis revealed that the organoids upregulated expression of key genes involved in spermatogenesis (Figure 2M). Expression of DAZL, a master translational regulator vital to SSC self-renewal and differentiation,[65, 66] was upregulated 9.7±1.6-fold. Accordingly, the quiescent SSC markers UTF1 and ID4[2] also increased 46.7± 8.8-fold and 5.8± 0.6-fold, as did KIT[2, 55], a marker of mitotic differentiating SSCs, by 9.4± 6.7-fold. DMRT1 and STRA8, two retinoic acid-stimulated genes responsible for regulating SSC entry into differentiation,[49, 67-70] were downregulated 18.9±14.9-fold and 24.4± 29.4-fold, demonstrating an improved resistance to retinoic acid stimulation over monolayer SSC cultures. Differentiating germ cells reduce their genetic content by half to become haploid spermatids, and to this end they undergo a specialized cell division process termed meiosis, wherein a cell divides twice to generate four genetically different daughter cells, each containing a single set of chromosomes. SYCP3, a marker of meiosis,[2] was upregulated in the organoids 7.1±0.7-fold. After completion of meiosis, haploid spermatid cells undergo morphological changes as they mature into final spermatozoa form, characterized by the development of an organelle with egg-penetrating enzymes called an acrosome, head compaction, and elongation of a flagellum structure. ZPBP expression is required for acrosome development,[71] and was seen to increase 160.8± 16.0-fold, while TNP1 and PRM2, two of the genes responsible for head compaction[72], increased 5.8±0.7-fold and 382.4± 17.9-fold.

Sectioning and H&E staining of the organoids revealed tubular internal structures reminiscent of native testicular seminiferous tubule architecture (Figure 2K-L). Immunostaining revealed the tubular structures to be mainly composed of GATA1^+^/SOX9^+^ hSCs and CD34^+^ hECs (Figure 2A-C), while ACTA2^+^ hPTMs and INSL3^+^ hLCs were located near the outer edges of the organoids (Figure 2E-F). Interestingly, the mature Leydig cell marker INSL3 and the mature Sertoli cell marker GATA1 were both highly expressed, indicating that the organoid system had the effect of maturing the hLCs and hSCs. VWF remained negative, suggesting that the hECs did not mature (Figure 2D). SYCP3^+^ meiotic stage differentiating germ cells, and a small number of PRM2^+^ post-meiotic stage differentiating germ cells, were noted within the organoids, in agreement with the RT-qPCR analysis (Figure 2G-H).

## Discussion

In this work we derived testicular somatic cells and SSCs from hiPSCs and showed that they spontaneously self-organize into 3-dimensional testicular-like tissues under the right conditions. After 12 days of culture in medium supplemented with hormones and growth factors to mimic the testicular *in vivo* microenvironment, the organoids matured and began spermatogenesis.

The ability of hiPSC-derived testicular organoids to adopt mature testicular cell phenotypes suggests their use as a surrogate cellular resource for establishing *in vitro* systems to mature primary human pre-pubertal testicular tissues. Cancer treatments are often gonadotoxic, therefore pre-pubertal patients are given the option of banking their pre-treatment testis tissues[73, 74] however maturing these tissues to generate spermatozoa for *in vitro* fertilization (IVF) therapies remains in the experimental stages, and is currently dependent upon extremely rare pre-pubertal testicular tissues with which to establish *in vitro* culture systems.[75-77] hiPSC-derived testicular organoids could serve as a suitable alternative resource to optimize such *in vitro* systems until larger banks of pre-pubertal tissues are available to researchers.

A second important finding was the ability of the hiPSC-derived organoid system to regulate spermatogenesis. *In vivo*, germ cells exist as a highly heterogeneous population, composed of quiescent, self-renewing and differentiating fractions,[2] and this balance is achieved by short range regulatory paracrine signaling from somatic cells. Self-renewal is stimulated by GDNF signaling,[78-81] while entry into differentiation is controlled by retinoic acid, whose bioavailability is regulated by Sertoli cell-secreted degradative enzymes.[82] Genetic markers for quiescent and self-renewing SSC populations were upregulated in the organoids in parallel with genetic markers of meiotic and post-meiotic stage differentiating germ cells, mirroring *in vivo*-like SSC heterogeneity suggestive of paracrine regulation within the organoids. Indeed, paracrine signaling could explain the noted decrease in retinoic acid-stimulated STRA8 expression in the organoids despite substantially increasing retinoic acid supplementation in the medium. This is in keeping with primary SSC studies showing that SSC resistance to retinoic acid stimulation is contingent on the topography of testis tissue facilitating short range paracrine signaling between specific cell types.[20, 21, 51]

This work illustrates the ability of hiPSCs to generate functional testicular organoids for the first time. We show that testicular somatic cells and SSCs derived from hiPSCs spontaneously organize into miniature tissues that resemble the microarchitecture of native testicular tissue, triggering somatic cell maturation and spermatogenesis. These organoids can serve as a limitless and accessible resource for the study of human testicular tissues *in vitro*.

## Materials and methods

### hiPSC cell culture

The hiPSC line used was the 1-DL-01 line from WiCell (hPSCRegID WISCi003-A).[83] hiPSCs were expanded on Growth Factor Reduced Matrigel (Corning, 354230) in mTeSR™-Plus medium (STEMCELL Technologies, 100-0276), at 37°C and 5% CO_2_, and passaged when 90% confluent using ReLeSR™ selective passaging reagent (STEMCELL Technologies, 05872) to maintain purity.

### Human ethics

Experiments using hiPSCs in this study were not subject to ethics approval from the University of British Columbia Research Ethics Boards or Stem Cell Oversight Committee, since they were derived from somatic cells and not intended for transfer into humans or non-human animals.

### Differentiation of Leydig cells from hiPSCs

Leydig cells (hLCs) were differentiated using a previously described method,[28] with a modification. The original protocol yielded 41.5% Leydig cells, likely because hiPSCs were spontaneously differentiated for 2 days prior to directed differentiation, allowing for endodermal and ectodermal lineage acquisition in addition to mesenchymal. To avoid this, we first directed the hiPSCs towards an intermediate mesenchymal fate using a previously described protocol,[84] wherein cells are treated with 5 µM CHIR99021 (STEMCELL Technologies, 72052) in Roswell Park Memorial Institute Medium 1640 (RPMI 1640, Gibco, 11875-093) for 36 hours, and then 1 µM all-trans retinoic acid (RA, STEMCELL Technologies, 72262) and 100ng/mL human recombinant Fibroblast Growth Factor 2 (FGF2, STEMCELL Technologies, 78003.1) for 2 days. These intermediate mesenchymal cells were then subjected to the protocol for Leydig cell differentiation. Cells were cultured throughout on Matrigel substrates in Dulbecco’s Modified Eagle Medium/Nutrient Mixture F-12 with 15mM HEPES (DMEM/F12, STEMCELL Technologies, 36254) 1X Insulin Transferrin Selenium Liquid Media Supplement (ITS, Millipore Sigma, I13146), 1% penicillin/streptomycin (Sigma Aldrich, P4333)and 1X GlutaMAX (Thermofisher, 35050079), 1% bovine serum albumin (BSA, Miltenyi Biotec, 130-091-376) and 5 ng/mL luteinizing hormone from human pituitary (LH, Sigma, L6420), with media changes every other day. From 0–7 days, 0.2 μM Smoothened Agonist (SAG, STEMCELL Technologies, 73412), 5 μM 22R-hydroxycholesterol (22R-OHC, Sigma, H9384), and 5 mM lithium chloride (Sigma, 62476) were added. From 7–10 days, 5 ng/mL human recombinant Platelet-Derived Growth Factor-AA (PDGF-AA, Peprotech, 100-13A) and 5 ng/mL FGF2 were added. From 10–17 days, 5 ng/mL PDGF-AA, 5 nM Insulin-Like Growth Factor 1 (IGF1, Peprotech, 100-11), and 10 μM testosterone (Toronto Research Chemicals, T155010) were added. From days 17–20, 10 ng/mL PDGF-AA and 10 ng/mL FGF2 were added. From days 20–25, 5 ng/mL LH (total of 10 ng/mL), 0.5 mM RA and 1 mM 8-bromo-cAMP (Peprotech, 2354843) were added. On day 25 cells the cells were switched to Leydig Cell Media (Sciencell Research Laboratories, 4511) for expansion, and passaged onto poly-L-lysine (PLL, Sciencell Research Laboratories, 0413) using TrypLE™ Express Enzyme (Thermofisher, 12604013).

### Differentiation of peritubular myoid cells from hipscs

hiPSC-derived peritubular myoid cells (hPTMs) were differentiated using the above protocol for hiPSC-derived Leydig cells with the modification that PDGF-AA is replaced by human recombinant Platelet-Derived Growth Factor-BB, as previously described [manuscript submitted] (PDGF-BB, Peprotech, 100-14B). After differentiation, hPTMs were expanded on PLL-coated plates in DMEM/F12, 1X ITS, 1% penicillin/streptomycin, 2.5% fetal bovine serum (FBS, Gibco, 12483-020), 10 ng/mL FGF2, 1 ng/mL human recombinant Leukemia Inhibitory Factor (LIF, STEMCELL Technologies, 78055.1) and 10 ng/mL human recombinant Epidermal Growth Factor (EGF, STEMCELL Technologies, 78006.1), and passaged using TrypLE™ Express Enzyme.

### Differentiation of Sertoli cells from hiPSCs

hiPSC-derived Sertoli cells (hSCs) were differentiated as previously described,[27] but with Matrigel-coated plates instead of feeder cells. hiPSCs were made into embryoid bodies using AggreWell™800 microwells (STEMCELL Technologies, 34811) in AggreWell™ EB Formation Medium (STEMCELL Technologies, 05893), supplemented with 50 ng/mL FGF2 and 30 ng/mL animal-free recombinant human Bone Morphogenic Protein 4 (BMP4, Peprotech, AF-120-05ET) for 24 hours. Embryoid bodies were then transferred to Matrigel-coated plates in DMEM/F12, 1X ITS, 1% penicillin/streptomycin, 1X GlutaMAX, 50 ng/mL recombinant human Fibroblast Growth Factor 9 (FGF9, Peprotech, 100-23), 500 ng/mL Prostaglandin D2 (PGD2, Peprotech 4150768), and 40 ng/mL animal component-free human recombinant Activin A (STEMCELL Technologies, 78132) for 12 days with media changes every other day. After 12 days the cells were switched to Sertoli Cell Medium (Sciencell, 4521) for expansion, and passaged onto PLL-coated plates using TrypLE™ Express Enzyme.

### Differentiation of endothelial cells from hiPSCs

hiPSC-derived endothelial cells (hECs) were differentiated as previously described.[85] hiPSCs were subjected to 5 μM CHIR99021 in STEMdiff APEL 2 Medium (STEMCELL Technologies, 05270) for 24 hours, followed by 25 ng/mL BMP4 for 24 hours, and 25 ng/mL BMP4 with 50 ng/mL human recombinant Vascular Endothelial Growth Factor 165 (VEGF-165, STEMCELL Technologies, 78073) for 2 days. On day 4, 6 million cells were collected and sorted with magnetic activated sorting (MACS), using a PE-conjugated antibody for anti-CD34, the EasySep™ Human PE Positive Selection Kit II (STEMCELL Technologies, 17664), and EasySep™ Magnet (STEMCELL Technologies, 18000). The purified cells were then expanded on PLL-coated plates in Endothelial Cell Growth Medium (Promocell, C-22010) and passaged using TrypLE™ Express Enzyme.

### Differentiation of SSCs from hiPSCs

hiPSC-derived SSCs (hSSCs) were differentiated as previously described.[30] hiPSCs were grown on Matrigel in Minimum Essential Medium Alpha (αMEM, Gibco A10490-01), 1X GlutaMAX, 1X ITS, 0.2% BSA, 1% penicillin/streptomycin, 1 ng/mL FGF2, 20 ng/mL animal component-free human recombinant Glial Cell Derived Neurotrophic Factor (GDNF, STEMCELL Technologies, 78139), 0.2% Chemically Defined Lipid Concentrate (Thermofisher, 11905031), and 200 µg/mL L-ascorbic acid (Sigma, A4544). Medium was changed every 2 days for 12-15 days. Cells were then switched to expansion medium as previously described, composed of StemPro-34 SFM (Thermofisher, 10639011) supplemented with 1X ITS, 30 μg/ml sodium pyruvate (Sigma, S8636), 1 μl/ml sodium DL-lactic acid solution (Sigma, L4263), 5 mg/ml BSA, 1% FBS, 1X GlutaMAX, 5 × 10^−5^ M 2-mercaptoethanol (Gibco, 31350-010), 1X Minimal Essential Medium (MEM) Vitamin Solution (Gibco, 11120052), 10^−4^ M ascorbic acid, 10 μg/ml biotin (Sigma, B4639), 30 ng/ml β-estradiol (Sigma, E2758), 60 ng/ml progesterone (Sigma, P8783), 20 ng/mL EGF, 10 ng/mL LIF, and 10 ng/mL GDNF. Cells were passaged onto CellAdhere™ Laminin-521 coated plates (STEMCELL Technologies, 77004) using TrypLE™ Express Enzyme (Gibco). 12 million cells were then purified for CD90^+^ cells using a PE-conjugated antibody for anti-CD90 (R&D Systems, FAB2067P), the EasySep™ Human PE Positive Selection Kit II and EasySep™ Magnet.

### Generation of organoids

Testicular organoids were made by mixing hLCs, hSCs, hSSCs, hPTMs and hECs at a 1:2:1:1:1 ratio and placing them into Aggrewell™800 microwells overnight. The ratio of hSC to hSSC in the organoids was chosen to be double that observed *in vivo* (4:1 SC:SSC),[86] to ensure an adequate pool of hSCs, while the remaining somatic cell ratios were approximated by observations of H&E stained human testicular biopsies. The following day the organoids were transferred to Ultra-Low Adherent Plates for Suspension Culture (STEMCELL Technologies, 38071), at 1 Aggrewell™800 well per well (of a 6-well plate), or roughly 300 organoids per well. We cultured organoids throughout in StemPro™-34 SFM media supplemented with 1X ITS, 30 μg/ml sodium pyruvate, 1 μl/ml DL-lactic acid solution, 5 mg/ml BSA, 1% FBS, 1X GlutaMAX, 5 × 10^−5^ M 2-mercaptoethanol, 1X MEM Vitamin Solution, 10^−4^ M L-ascorbic acid, 10 μg/ml biotin, 30 ng/ml β-estradiol, 60 ng/ml progesterone, 10 ng/mL LIF, 10 ng/mL EGF, 100 ng/mL Follicle Stimulating Hormone from human pituitary (FSH, Millipore Sigma, F4021), 10 ng/mL LH, and 1 µM metribolone (Toronto Research Chemicals, M338820), 100 ng/mL BMP4, 100 ng/mL animal-free recombinant human Stem Cell Factor (SCF, Peprotech, AF-300-07), and 10 µM RA. Each well constituted a single biological replicate in our study.

### RT-qPCR

A sample of ∼50 organoids was taken from each biological replicate and RNA was extracted using an RNeasy Plus Micro Kit (Qiagen, 74034), and checked for integrity using an Agilent 2200 Tapestation System with High Sensitivity RNA Screentape (Agilent, 5067-5579), High Sensitivity RNA ScreenTape Sample Buffer (Agilent, 5067-5580), and High Sensitivity RNA ScreenTape Ladder (Agilent, 5067-5581). cDNA was generated using iScript™ Reverse Transcription Supermix (Bio-Rad, 1708840), followed by pre-amplification with SsoAdvanced™ PreAmp Supermix (Bio-Rad, 1725160) and pre-amplification primers with a Bio-Rad Tetrad2 Peltier Thermal Cycler. RT-qPCR was done with SsoAdvanced™ Universal SYBR® Green Supermix (Bio-Rad, 1725270) on a LightCycler96 (Roche). Pre-amplification primers and regular primers used were PrimePCR™ SYBR®® Green Assays (Bio-Rad) as listed in **Table 1**. Technical replicates were carried out in triplicate. Analyses was done in Excel and GraphPad Prism Software. Ct values were normalized to Glyceraldehyde-3-Phosphate Dehydrogenase (GAPDH). Technical replicate outliers were detected using Grubbs’ Test, with α=0.05. Results of RT-qPCR are presented as the average Relative Quantification (RQ=2^-ΔΔCt^) values and standard deviations of the biological replicates. Any undetected samples were given a Ct value of the maximum detected cycles plus 1.

**Table 1.**
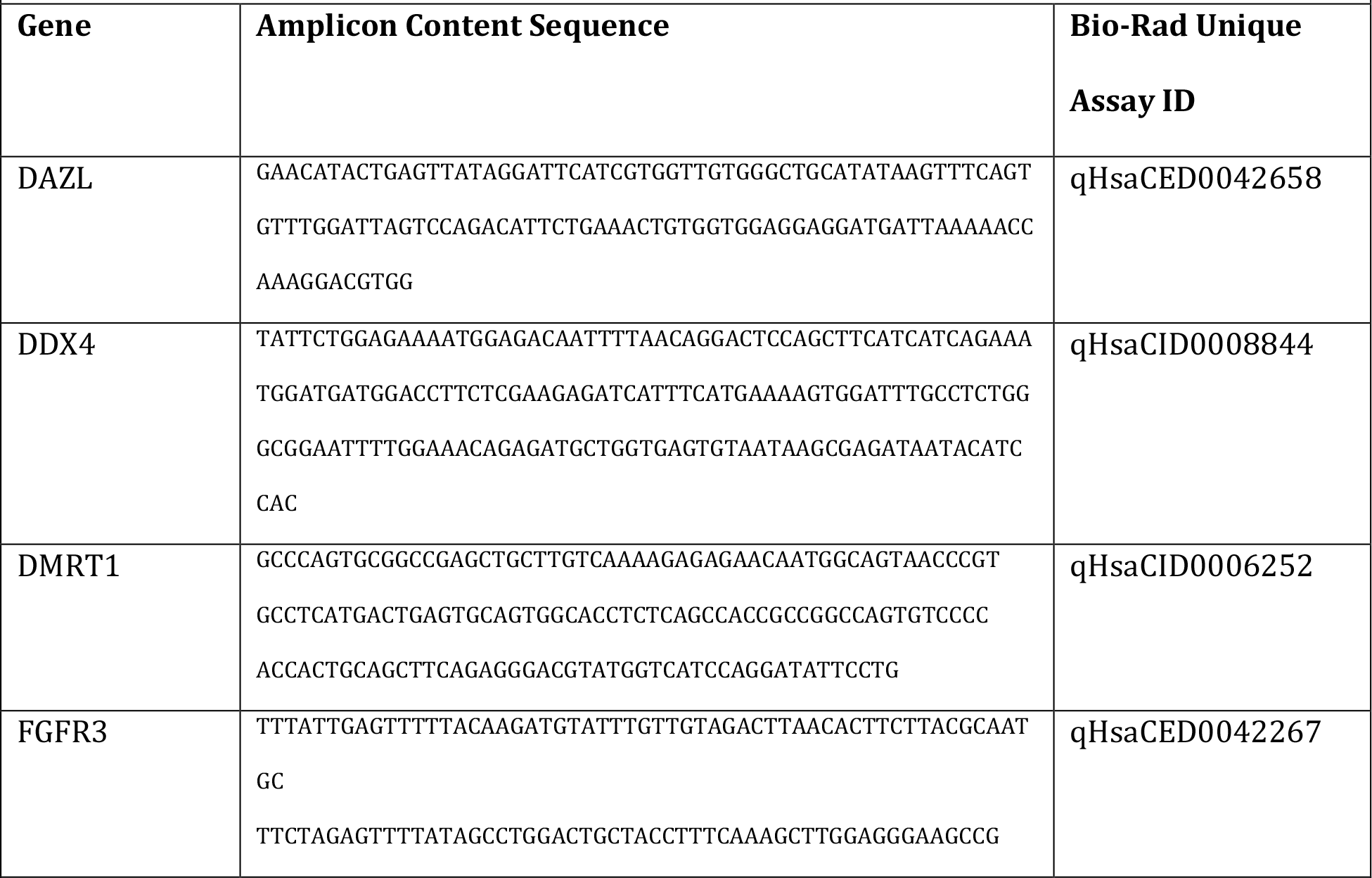
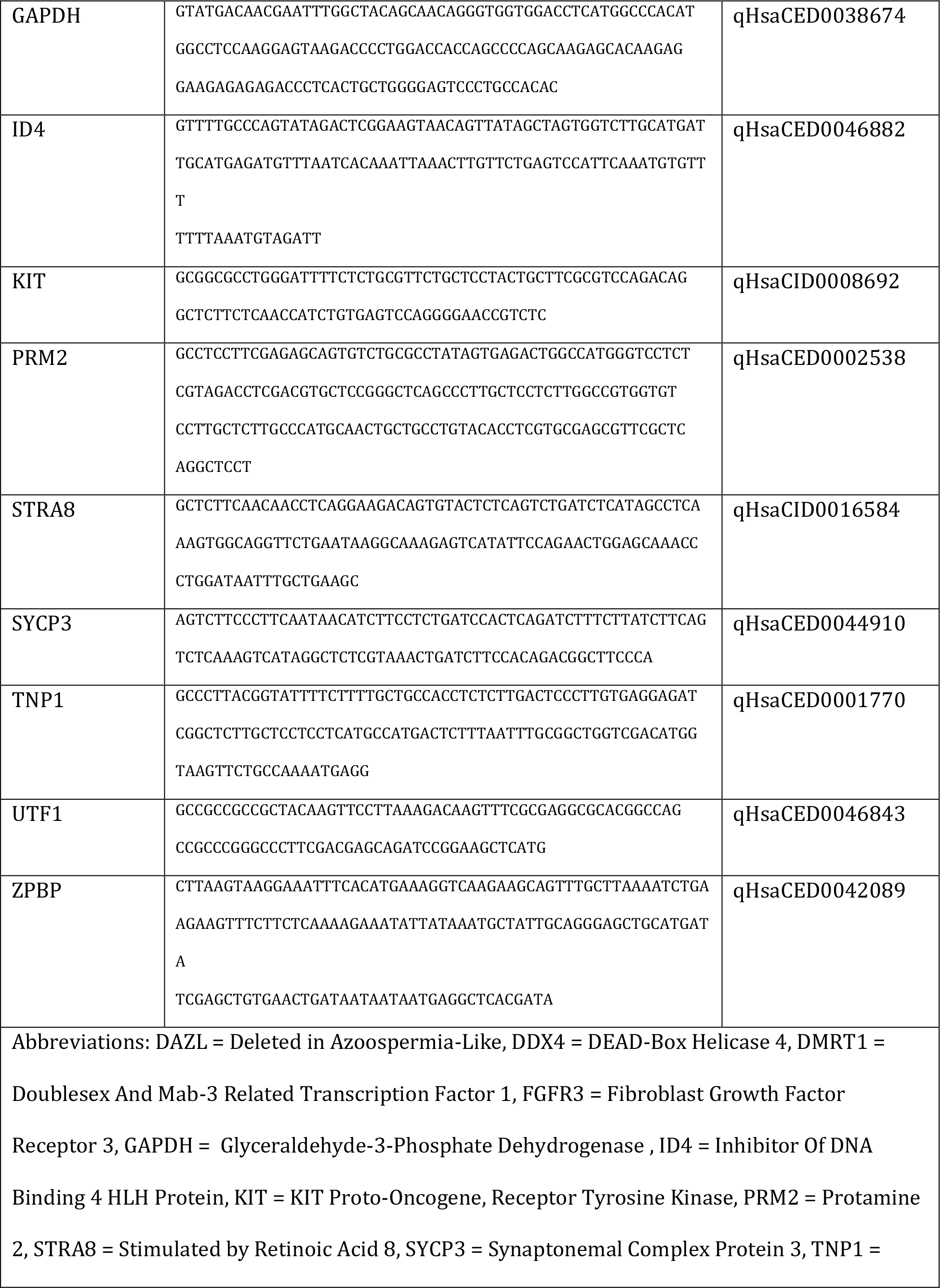
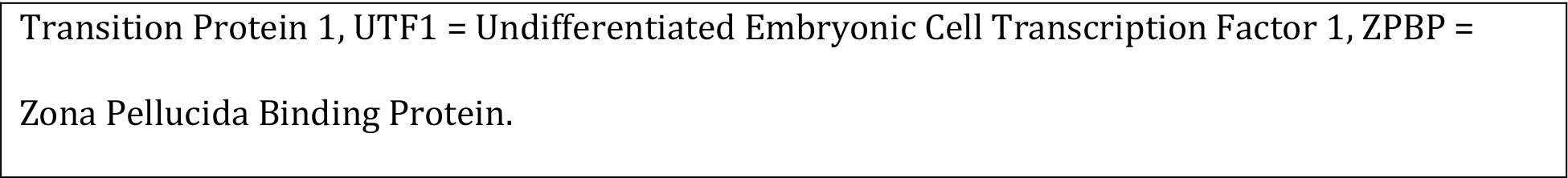
Amplicon content sequences and Unique Assay IDs for Bio-Rad PrimePCR™ primer pairs used in real time quantitative polymerase chain reaction (RT-qPCR) assays. Amplicon content sequences (amplicon sequence with additional base pairs added to the beginning and/or end of the sequence), and Unique Assay IDs for Bio-Rad PrimePCR™ primer pairs used.

### Immunocytochemistry

Cells were fixed for 15 minutes in 4% paraformaldehyde solution (PFA, Thermo Scientific, J19943-K2), permeabilized for 15 minutes in 0.1% Triton X-100 (Sigma, X100) in phosphate buffered saline (PBS), and blocked for 2 hours in 5% normal goat serum (NGS, Abcam, ab7481) in PBS. Primary antibodies were diluted in PBS as follows: anti-Hydroxy-Delta-5-Steroid Dehydrogenase, 3 Beta- And Steroid Delta-Isomerase 1 (HSD3β, Novus Biologicals, NB110-7844) 1:200, anti-Myosin Heavy Chain 11 (MYH11, Abcam, ab212660) 1:1000, anti-Octamer Binding-Protein 4 (OCT4, Abcam, ab184665) 1:500, anti-SRY-Box Transcription Factor 9 (SOX9, Abcam, ab76997) 1:500, anti-GATA Binding Protein 4 (GATA4, Abcam, ab84593) 1:100, anti-Nestin (Millipore, MAB5326) 1:200, anti-Thy-1 Cell Surface Antigen (THY1/CD90, Abcam, ab133350) 1:200, anti-GATA Binding Protein 1 (GATA1, Abcam, ab28839) 1:300, anti-Insulin Like 3 (INSL3, Novus Biologicals, NBP1-81223) 1:500, anti-CD34 Molecule (CD34, Abcam, ab81289) 1:250, anti-Von Willebrand Factor (VWF, Invitrogen, MA5-14029) 1:100, anti-GDNF Family Receptor Alpha 1 (GFRA1, Abcam, ab84106) 1:200, anti-Stage-specific embryonic antigen-4 (SSEA4, Abcam ab16287) 1:300, anti-G Protein-Coupled Receptor 125 (GPR125, Abcam, ab51705) 1:200, anti-Vimentin (Abcam, ab20346) 1:1000, anti-STRA8 (Millipore ABN1656) 1:100, and incubated overnight at 4°C in the dark. Cells were rinsed 3 times with PBS for 15 minutes each at 4°C in the dark. Goat anti-Rabbit IgG (H+L) Highly Cross-Adsorbed Secondary Antibody Alexa Fluor 488 (Thermofisher, A-11034) or Goat anti-Mouse IgG (H+L) Highly Cross-Adsorbed Secondary Antibody Alexa Fluor 568 (Thermofisher, A-11031) were diluted 1:200 in PBS and incubated with the cells for 4 hours at 4°C in the dark. Cells were rinsed another 3 times with PBS for 15 minutes each at 4°C in the dark. 4′,6-diamidino-2-phenylindole (DAPI, Abcam, ab228549) was diluted to 2.5 µM in PBS and added to the cells for 15 minutes in the dark at room temperature, and then replaced by PBS. Cells were imaged using a Zeiss AXio Observer microscope equipped with laser excitation and fluorescence filters for AlexaFluor 488, AlexaFluor 568 and DAPI, and images were processed using ZEN Blue and ImageJ software.

### Immunofluorescence staining of 3-D organoids

Sectioning and immunofluorescent staining of the organoids was done as described previously,[87] with some modifications. The remaining ∼250 organoids from each biological replicate were fixed for 24 hours in 4% PFA solution, followed by 24 hours in 30% sucrose (Sigma, S1888) solution in PBS. The organoids were then re-suspended in 0.5 mL 7.5% gelatin from porcine skin (Sigma, G2500) with 10% sucrose in PBS and incubated at 37°C for 1 hour to allow the gelatin to penetrate the organoids before they were gently pipetted into 10×10×5mm cryomolds (Tissue-Tek, 4565) and flash frozen. Flash frozen samples were then cut into 6 µm sections using a cryostat with the block temperature set to −30°C, and placed onto Superfrost Plus microscope slides (Fisherbrand, 22-037-246). A hydrophobic barrier was drawn around the tissue sections using a hydrophobic PAP pen (Abcam, ab2601), and the sections were incubated with 0.1% Tween 20 (Sigma, P1379) in PBS at 37°C for 10 minutes to dissolve the gelatin solution. Sections were blocked for 1 hour with 5% NGS in 0.1% Tween 20 in PBS in a humidity chamber at room temperature. Primary antibodies were diluted in 0.1% Tween 20 in PBS with 0.05% sodium azide and 5% Bovine Serum Albumin (BSA) as follows: 1:300 anti-GATA1, 1:500 anti-Actin Alpha 2, Smooth Muscle (ACTA2, Thermofisher, 14-9760-82), 1:500 anti-INSL3, and 1:300 anti-SYCP3 (Novus Biologicals, NB300-232). Sections were incubated in the diluted primary antibodies overnight in a humidity chamber at room temperature. The following day the sections were rinsed 3 times with 0.1% Tween 20 in PBS for 30 minutes each in a humidity chamber, followed by incubation for 2 hours in Goat anti-Mouse IgG (H+L) Cross-Adsorbed Secondary Antibody Alexa Fluor 647 (Thermofisher, A-21235) or Goat anti-Rabbit IgG (H+L) Cross-Adsorbed Secondary Antibody Alexa Fluor 647 (Thermofisher, A27040) diluted 1:2000 in 0.1% Tween 20 in PBS. We used only Alexa Fluor 647 because we noted the organoids to auto-fluoresce in the 488 nm and 568 nm wavelengths, but not in the 647 nm wavelength. The sections were then rinsed 3 more times for 30 minutes each with 0.1% Tween 20 in PBS, with the final rinse containing 2.5 µM DAPI. Sections were imaged immediately using a Zeiss AXio Observer microscope equipped with laser excitation and fluorescence filters for AlexaFluor 647 and DAPI, and images were processed using ZEN Blue and ImageJ software.

### Hematoxlyin and Eosin (H&E) staining

Embedding and sectioning was completed as described in Immunofluorescence staining. After sectioning, a hydrophobic barrier was drawn around the tissue sections using a hydrophobic PAP pen, and they were incubated with 0.1% Tween 20 in PBS for 10 minutes at 37°C to dissolve the gelatin solution. The slides were then placed in hematoxylin solution (Sigma Aldrich, MHS32) for 3 minutes, followed by tap water for 1 minute 2 times in separate containers, 0.1% sodium bicarbonate (Sigma S5761) solution for 1 minute, tap water for 1 minute, 95% ethanol for 1 minute, Eosin Y reagent (Sigma Aldrich, HT110216) for 1 minute, 95% ethanol for 1 minute, 100% ethanol for 1 minute 2 times in separate containers, and 3 minutes in xylene (Fisher Scientific X5-500). 1-2 drops of CytoSeal mounting media (Thermofisher, 8312-4) was placed on each slide, a coverslip was placed on top, and the slides were allowed to dry overnight in a fume hood. The organoids were imaged using an Olympus BX-UCB from PerkinElmer Life Sciences and VECTRA system.

### Statistical analysis

Statistics were performed using GraphPad Prism software. Each experiment was performed in biological triplicate, and in the case of RT-qPCR, in technical triplicate as well. Significance between groups was determined by comparing ΔCt values using a student’s unpaired two-tailed t-test, with α=0.05. The standard deviations between biological replicates are represented by error bars in the figures, and statistically significant differences between groups is denoted by * in the figures.

## Glossary

Differentiating: A term for the process of stem cell development into specialized cell types.
endothelial cells: Endothelial cells form the barrier between vessels and tissue and control the flow of substances and fluid into and out of a tissue.
Gonadotoxic: Gonadotoxicity is the temporary or permanent damage to ovaries or testes after exposure to certain substances or drugs
haploid: Haploid is the quality of a cell having a single set of chromosomes.
human induced pluripotent stem cells: iPSC are derived from skin or blood cells that have been reprogrammed back into an embryonic-like pluripotent state that enables the development of an unlimited source of any type of human cell needed for therapeutic purposes.
in vitro: Outside of the body, typically referring to a laboratory setting
in vitro fertilization (IVF): In vitro fertilization (IVF) is a method of assisted reproduction that involves the removal of eggs from the body, and the combining of eggs and sperm in the embryology laboratory to form embryos. The resulting embryos can then be placed into the uterus in the hopes of achieving a pregnancy.
in vivo: Within the body
Leydig cells: Leydig cells, also known as interstitial cells of Leydig, are found adjacent to the seminiferous tubules in the testicle. They produce testosterone in the presence of luteinizing hormone (LH)
Matrigel: Matrigel is the trade name for a gelatinous protein mixture secreted by Engelbreth-Holm-Swarm (EHS) mouse sarcoma cells produced by Corning Life Sciences.
meiosis: Meiosis is a process where a single cell divides twice to produce four cells containing half the original amount of genetic information.
organoid: An organoid is a miniaturized and simplified version of an organ produced in vitro in three dimensions that shows realistic micro-anatomy. They are derived from one or a few cells from a tissue, embryonic stem cells or induced pluripotent stem cells, which can self-organize in three-dimensional culture owing to their self-renewal and differentiation capacities.
paracrine signaling: Paracrine signaling is a form of cell signaling, a type of cellular communication in which a cell produces a signal to induce changes in nearby cells, altering the behaviour of those cells.
peritubular myoid cells: A peritubular myoid cell is one of the smooth muscle cells which surround the seminiferous tubules in the testis.
personalized medicine: Personalized medicine, also referred to as precision medicine, is a medical model that separates people into different groups— with medical decisions, practices, interventions and/or products being tailored to the individual patient based on their predicted response or risk of disease
quiescent: A term for “reserve” pools of stem cells that in a state of inactivity or dormancy.
Sertoli cells: A Sertoli cell (a kind of sustentacular cell) is a “nurse” cell of the testicles that is part of a seminiferous tubule and helps in the process of spermatogenesis, the production of sperm. It is activated by follicle-stimulating hormone (FSH) secreted by the adenohypophysis, and has FSH receptor on its membranes
spermatid: The spermatid is the haploid male gametid that results from division of secondary spermatocytes.
spermatogenesis: Spermatogenesis is the process by which haploid spermatozoa develop from spermatogonial stem cells in the seminiferous tubules of the testis
spermatogonial stem cells: A spermatogonial stem cell, also known as a type A spermatogonium, is a spermatogonium that does not differentiate into a sperm cell. Instead, they continue dividing into other spermatogonia or remain dormant to maintain a reserve of spermatogonia.
tissue banking: The process of cryopreserving tissues for long-term storage and viability.

## Acknowledgements

The authors would like to acknowledge the Vancouver Prostate Centre for their financial support, and the Lange Lab at the Vancouver Prostate Centre for their technical support.

